# Generation of Cynomolgus Monkeys (*Macaca fascicularis*) Embryos with ICSI Based on the MII-Stage Oocytes Acquired by Personalized Superovulation Protocol

**DOI:** 10.1101/475897

**Authors:** Zhangqiong Huang, Yun Li, Qinfang Jiang, Yixuan Wang, Kaili Ma, Qihan Li

## Abstract

The use of cynomolgus monkeys (*Macaca fascicularis*) has facilitated the establishment of gene-modified animal models, and the appropriate number of high-quality mature oocytes (MII-stage oocytes) are crucial in assisted reproductive technologies (ART) of cynomolgus macaques. In this study we performed 3 different superovulation protocols on 24 female cynomolgus monkeys aimed to improve the proportion of MII-stage oocyte retrieval. The results suggested some differences in the proportion of MII-stage oocytes between the personalized superovulation protocol and the superovulation protocols I and II. Afterwards, we performed Intracytoplasmic sperm injection (ICSI) on 22 MII-stage oocytes derived from one animal with personalized superovulation protocol, obtained 15 embryos of 2-4-cells, 2 cases of successful pregnancy after transferring to 4 surrogate female, and obtained 3 aborted fetuses. These results suggested that personalized superovulation protocol incorporated the menstrual cycle length of each cynomolgus monkey, and the timing of oocytes retrieval based on the equation: menstrual cycle length/2 ± 1, which increased the rate of MII-stage oocytes acquired and generation of cynomolgus monkeys embryos with ICSI based on these oocytes, and finally could successfully develop into fetus under conditions that effectively guarantee the synchronization of the surrogate mothers.

**SUMMARY STATEMENT:** The personalized superovulation protocol based on the menstrual cycle length of each cynomolgus monkey can increase the rate of MII-stage oocytes acquired, and finally could successfully develop into fetus under conditions that minimize damage to the surrogate mothers and effectively guarantee the synchronization of the surrogate mothers.

## INTRODUCTION

Nonhuman primates have been widely used for biomedical research and as transgenic animal models for humans particularly because of their similarity to humans in genetics, endocrine processes, menstrual cycle physiology, and timing of prenatal development (Esch et al., 2008; Gibbs et al., 2007; Stouffer and Woodruff, 2017; Zuo et al., 2017). Rhesus monkeys (*Macaca mulatta*) and cynomolgus monkeys (*Macaca fascicularis*) are the most common representatives of nonhuman primates in terms of research. The first “test tube” rhesus monkey was born in 1984(Bavister et al., 1984), after that, other primate species include cynomolgus monkey(Balmaceda et al., 1988), pigtailed macaque(Kubisch et al., 2006), marmosets(Sasaki et al., 2009), bamboo(Simerly et al., 2010), and African green monkey(Shimozawa et al., 2010) have been established by assisted reproductive technologies (ART), However, due to the lack of resources, high cost, complex procedures as well as the long reproductive cycle and so on, the development speed and success rate of ART in monkeys are still far from those of rodents and humans. In particular, the use of ART in cynomolgus monkeys remains modest. Compared with rhesus monkeys, the breeding of cynomolgus monkeys is not subject to seasonal restrictions, enabling their increased use in transgenic animal models. Regardless of the animal model used, the quality of oocytes is crucial in ART, and the appropriate number of high-quality mature oocytes (MII-stage oocytes) has long been considered as a primary limiting factor for *in vitro* fertilization-embryo transfer (IVF-ET). Numerous studies have shown that the quality of mature oocytes *in vivo* exceeds the quality of matured oocytes via *in vitro* maturation (Dupont et al., 2009; El and Haaf, 2013; Ko et al., 2015; Schramm et al., 2003). Most importantly, the quality of the oocytes directly impacts the quality of the embryos and thus affects the rates of oocytes fertilization, embryonic implantation, and pregnancy (Alvarez et al., 2013; Dupont et al., 2009; El and Haaf, 2013; Ko et al., 2015; Schramm et al., 2003). Therefore, the high-quality embryos is critically dependent on the superovulation method based on obtaining high-quality mature oocytes, but pregnancy and fetus production depend on embryo transfer, and uterine–embryo synchrony.

In the past 20 years, despite a few previous reports on promoting ovulation in non-human primate, the average of all menstrual cycles was used to calculate the regimen and frequency of hormonal injection. That is, as each monkey’s ovulation time is considered to be the mean for all monkeys, the time interval between the injection of the first hormone and the collection of oocytes was identical for all monkeys. Therefore, the number and proportion of MII-stage oocytes obtained were quite different (Chen et al., 2011; Nyachieo et al., 2009; Stouffer, 2002). In general, the results have been unsatisfactory. It is difficult to determine a technology that can provide consistent and reliable results.

In the present study, a superovulation protocol personalized to each cynomolgus monkey’s menstrual cycle was used, with intervals of 35–36 h between the injection of human chorionic gonadotropins (hCG) and oocyte collection. This protocol was compared with typical superovulation protocols using an average menstrual cycle for all cynomolgus monkeys. Then generated embryos by intracytoplasmic sperm injection (ICSI) on 22 MII-stage oocytes obtained from one of the personalized superovulation monkeys, and these embryos were transferred into the oviduct of recipient female selected from three key points. The objective of this study was to obtain a higher proportion of MII-stage oocytes through different superovulation protocols, and verified that these MII-stage oocytes could successfully produce the fetus with minimizing the harm of the female surrogate cynomolgus monkey.

## RESULTS

### Oocyte retrieval

The retrieved oocytes were classified into maturational stages: (1) immature oocytes at the germinal vesicle (GV, intact germinal vesicle); (2) immature oocytes at the metaphase I status (MI, no germinal vesicle, no polar body); and (3) mature oocytes at the metaphase II status, (MII, with the first polar body). The mean number of oocytes harvested using the personalized superovulation protocol was 26.6 ± 9.0, compared with 16.4 ± 7.3 for the superovulation protocol I (*P* < 0.05) (Fig. 2F). The proportion of MII oocytes retrieved from the personalized superovulation protocol was 58.5% ± 9.9%, greater than that from either superovulation protocol I (33.2% ± 5.1%, *P* < 0.01) or II (21.5% ± 19.6%, *P* < 0.01) (Fig. 2F). The proportion of GV-stage oocytes from the personalized superovulation protocol was significantly less than that from either superovulation protocol I or II (*P* < 0.01 and *P* < 0.01, respectively) (Fig. 2F). With respect to a single monkey, no difference in the total number of oocytes collected was observed on comparing the first ovulation induction without GnRH-a and using superovulation protocol I with the second ovulation induction using the personalized superovulation protocol. However, the difference in the percentage of GV- and MII-stage oocytes obtained using superovulation protocol I and the personalized superovulation protocol was obvious (*P* < 0.05 and *P* < 0.05, respectively) (Fig. 2G). In addition, the proportion of MII-stage oocytes between the first ovulation induction using superovulation protocol II and the second induction using the personalized superovulation protocol was different (*P* < 0.05) (Fig. 2H). On the other hand, as evident from Fig. 2F and 2G, and 2H, oocyte progression from the GV stage to the MI and MII stages and the proportion of MII-stage oocytes from the personalized superovulation protocol relative to the superovulation protocols I and II showed a tendency to increase. However, this tendency was exactly the opposite in superovulation protocols I and II; and a decrease in GV-stage oocytes was observed in the personalized superovulation protocol.

**Fig. 1.**
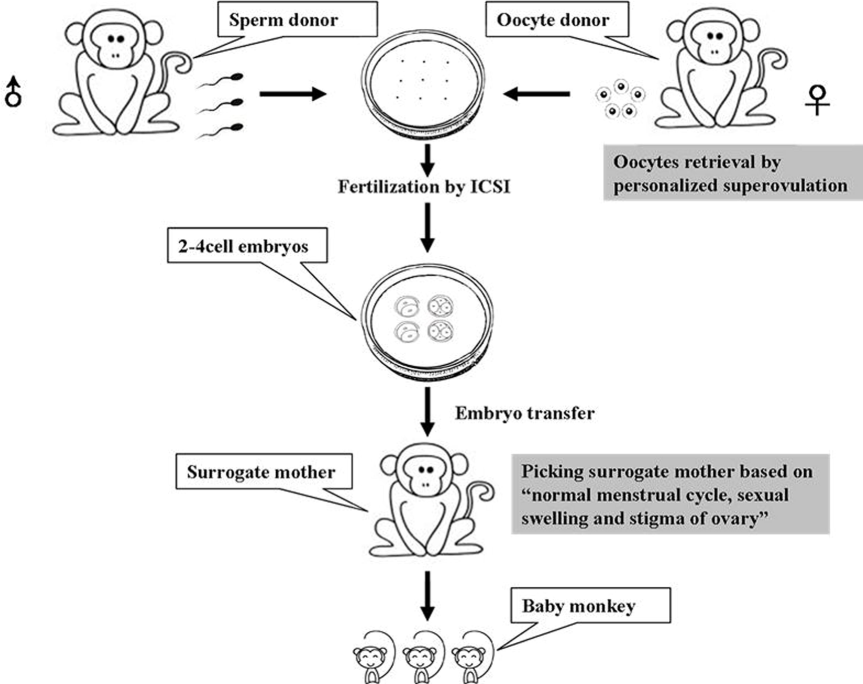
The highlights in superovulation and picking recipients in female cynomolgus monkeys in this study. Increasing the rate of MII-stage oocytes acquired by personalized superovulation protocol that incorporated the menstrual cycle length of each cynomolgus monkey. According to the “the normal menstrual cycle, sexual swelling and stigma of ovary”, the three points were highly efficient and accurate to ensure the synchronization of the surrogate mothers.

**Fig. 2.**
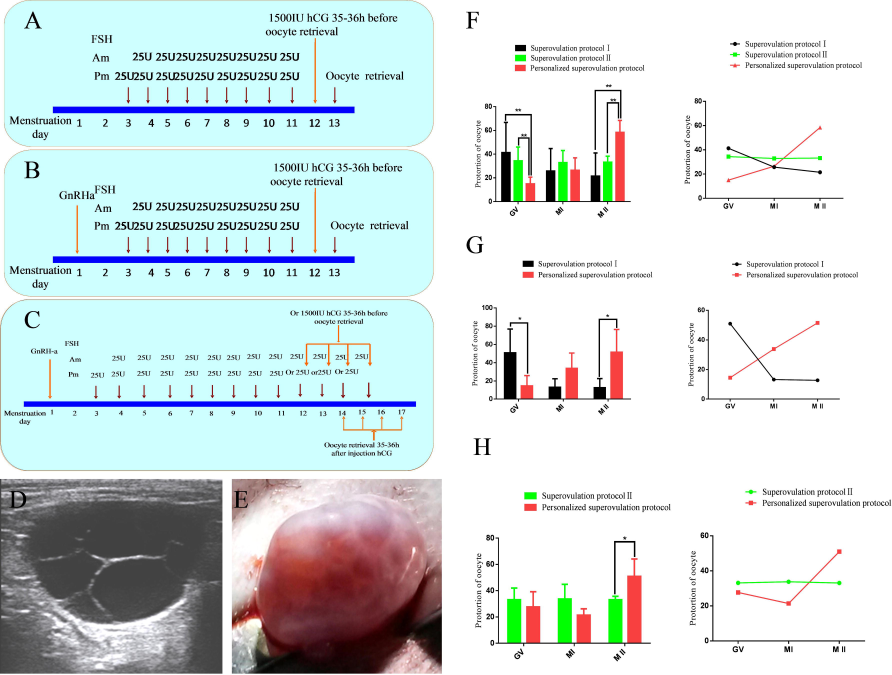
Superovulation protocol. (A) Superovulation protocol I. (B) Superovulation protocol II. (C) Personalized superovulation protocol. (D) B-type ultrasound images of the ovary and follicles after ovulation induction with personalized superovulation protocol. (E) Surgical removal of ovaries after ovulation induction with personalized superovulation protocol. (F) The total number and proportion of oocytes retrieved after ovulation induction. “*” indicates a difference(p ≤0.05), “**” and “##” indicates a significant difference(p ≤0.01). The results of superovulation protocol I and the personalized superovulation protocol showed that the average numbers of oocytes retrieved were 16.4 ± 7.3 and 26.6 ± 9.0, respectively (P<0.05). The proportion of MII-stage oocytes retrieved from the personalized superovulation protocol was 58.5%±9.9%, significantly greater than the 33.2% ± 5.1% from superovulation protocol I (P<0.01) or the 21.5% ± 19.6% from superovulation protocol II (P<0.01). The proportion of GV-stage oocytes from the personalized superovulation protocol were 15.0% ± 5.6%, significantly less than from either protocol I (41.3% ± 25.5%, P<0.01) or II (34.4% ± 11.6%, P<0.01). Oocyte progression from the GV stage to the MI and MII stages and the proportion of MII-stage oocytes from the personalized superovulation protocol relative to the superovulation protocols I and II showed a tendency to increase. However, this tendency was exactly the opposite in superovulation protocols I and II. (G) “*” indicates a difference, “**” indicates a significantly difference. Results in the same monkey between superovulation protocol I and the personalized superovulation protocol showed that the respective proportions of GV-stage oocytes were 51.0% ± 25.9% and 14.6% ± 11.3%, respectively (P<0.05). The proportion of MII-stage oocytes retrieved from the personalized superovulation protocol was 51.5% ± 24.9%, significantly higher than that from superovulation protocol I (12.8% ± 9.9%, P<0.05). The proportion of MII-stage oocytes from the personalized superovulation protocol relative to the superovulation protocols I showed a tendency to increase; and a decrease in GV-stage oocytes and elevation in MII-stage oocyte proportion were observed in the personalized superovulation protocol. (H) “*” indicates a difference. Results in the same monkey between superovulation protocol II and the personalized superovulation protocol showed that the proportions of MII-stage oocytes retrieved were 34.1% ± 2.7% and 51.0% ± 13.2%, respectively (P<0.05). The proportion of MII-stage oocytes from the personalized superovulation protocol relative to the superovulation protocols II showed a tendency to increase; and a decrease in GV-stage oocytes and elevation in MII-stage oocyte proportion were observed in the personalized superovulation protocol.

### Intracytoplasmic sperm injection

ICSI was performed on 22 MII-stage oocytes within 4 h post-isolation from one monkey that was based on personalized superovulaion method (Fig. 3A and 3B). Out of 22 MII-stage oocytes, 21 of them (95.4%) had an extruded second polar body(PB) after the injection and culture for 7-12 h, at last 20 of 22 (90.9%) occurred distinct pronuclei (Fig. 3C) within 12-18 h post - ICSI, 20-24 h after ICSI, 15 of 20 (75%) zygotes develop to 2-cell stage(Fig. 3D) and 3-cell stage; 4 embryos cleaved to the 3- to 4-cell stage within 26-28 h post - ICSI (Fig. 3E and 3F).

**Fig. 3.**
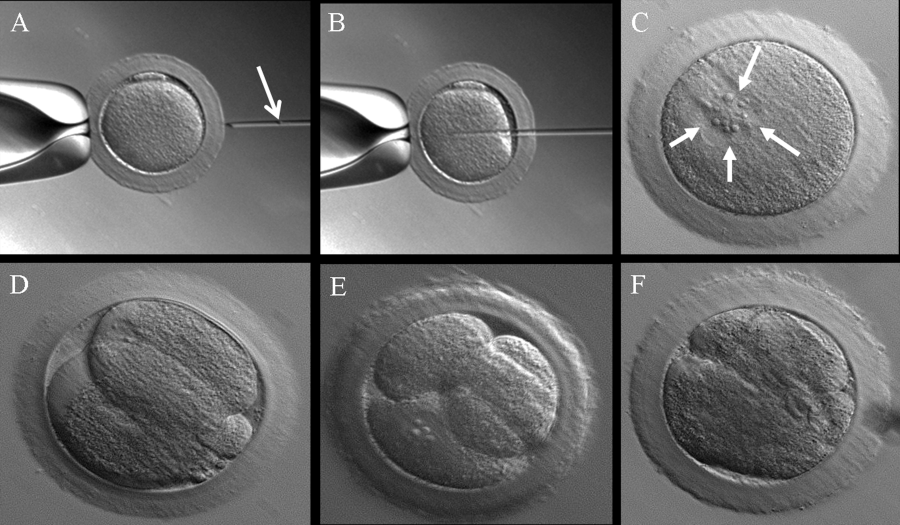
Intracytoplasmic sperm injection. (A) MII-stage oocytes are held so that the first polar body is positioned at 12 o’clock, and push a single sperm (aspirated tail first) to the tip of the injection pipet (arrow). (B) The injection pipet insert into oocyte cytoplasm through the zona pellucida from 3 o’clock. (C) zygotes occurred distinct pronuclein within 12-18 h post – ICSI, after several hours, the pronuclei were fused(arrow). (D) zygotes develop to 2-cell stage 20-24 h after ICSI. (E) zygotes develop to 3-cell stage 24-26 h after ICSI. (F) zygotes develop to 4-cell stage 26-28 h after ICSI.

### Embryo transfer

In order to verify whether MII-stage oocytes from the personalized ovulation protocol could develop into a fetus, 2-4-cell cleaved stage embryos were transferred into the oviducts of four female recipients for pregnancy establishment under laparoscopic guidance. There was stigma of ovulation on the ovary of female cynomolgus monkey recipients by observing the sexual swelling (Fig. 4F and 4G). Of these 11 embryos at 2-cell stage were transplanted to 3 female surrogates, and the other 4 embryos at 3- to 4-cell stage were transplanted to 1 female surrogate. Thirty days after transplantation, pregnancy was confirmed in 2 female recipients by ultrasound examination, 1 of them carried only gestational sacs; 1 of them with 3 fetuses (triplet) preterm births on the 103^rd^ day after embryo transfer, 1 of 3 was male, 2 of 3 was female (Fig. 5A); The rest surrogates were not pregnant.

**Fig. 4.**
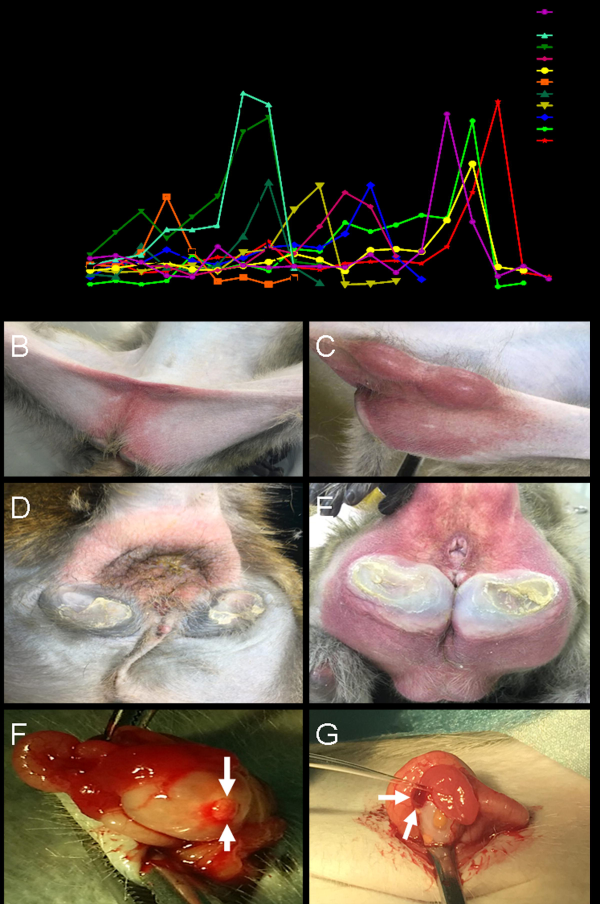
The characteristic of sexual skin of female cynomolgus monkey at two different stages of the menstrual cycle and ovary state of recipient female. (A) Estradiol detection: serum estradiol concentrations during days 7-20 after menstruation of female cynomolgus monkeys, one monkey first appeared E2 peak on the 7th day after menstruation, another peaked on day 20 after menstruation. (B) Non-ovulation characteristics of young female, turgescent area obviously narrowing and color decrease. (C) Ovulation characteristics of young female, in the early stage of sexual maturity, the vaginal area is swollen, reddened and subcutaneous groin subcutaneously forms two saclike protrusions of testicular size as the main features, and it has a smooth, shiny appearance and is deep, intense red color, periodic changes, and then did not appear. (D) Non-ovulation characteristics of adult female: turgescent area obviously narrowing, limit to reproductive anal area and tail root, and the labia and clitoris have many wrinkles with an overall pinkish red color. (E) Ovulation characteristics of adult female: turgescent area extended to anus and thighs, the perineal skin is fully distended with no wrinkles and smooth and most intense bright red color. (F) An new stigma (arrow) above the ovary of the recipient female. (G) Ovary with an new stigma (arrow), then 2–4-cell-stage embryos were transferred into the oviduct of a recipient female through the tubal fimbria. Bar = 15-20 mm.

**Fig. 5.**
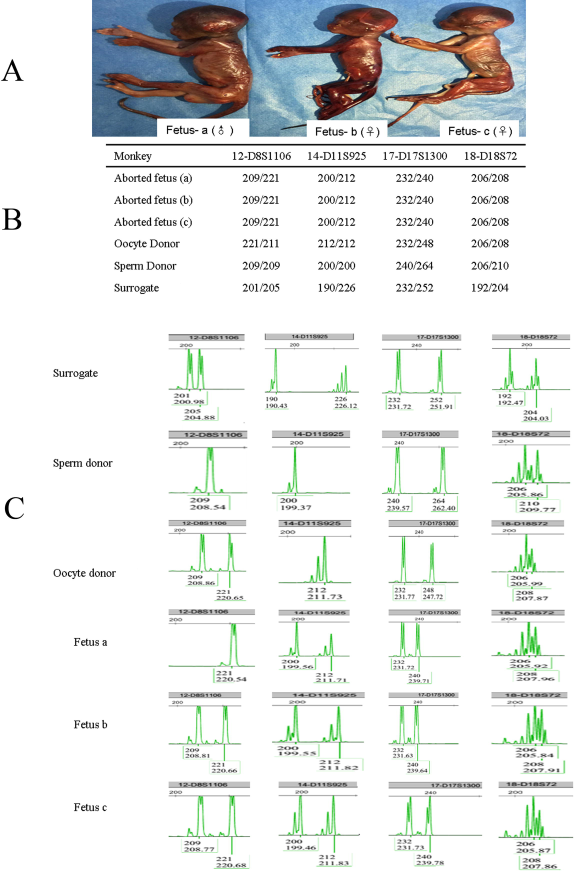
Parentage verification of aborted fetus. (A) Images of aborted fetus a, b and c. Three fetuses generated based on the MII-Stage oocytes acquired by personalized superovulation protocol but spontaneously aborted on day 103 post- embryos transfer. (B) Four examples of short tandem repeats (STRs) from fetus a, b,c, oocyte donor, sperm donor and surrogate, half of them were from the oocyte donor and half from the sperm donor. Complete list of STRs are shown in Data 1. (C) Examples of electrophoreses signal about amplification products of D8S1106, D11S925, D 17S1300, D18S72 sites for fetus a, b, c, oocyte donor, sperm donor and surrogate, showing fetus a, b and c have parent-child relationship with oocyte donor and sperm donor.

### Parentage verification of aborted fetus

STRs (short tandem repeats) DNA analyses was used by Liu and Tachibana et al. to detect DNA from cloned cynomolgus monkey by somatic cell nuclear transfer (SCNT)(Liu et al., 2018a; Tachibana et al., 2009). We performed STRs analysis on oocyte donor, sperm donor, surrogate and aborted fetuses using 20 microsatellite DNA loci, DNA from the hind limbs of aborted fetuses b and c, half of the 20 STRs were from the oocyte donor and half from the sperm donor. Half of the D12S364 locus of fetus a was from the sperm donor, and the other half may not show a genetic relationship with the oocyte donor due to mutation or deletion, but the other 19 STRs loci were half from the sperm donor and half from the oocyte donor. These genetic analyses confirmed the fetus a, b and c have parent-child relationship with oocyte donor and sperm donor. Six animals STRs were shown in Fig. 5C and complete information on 20 STRs loci were shown in Data1.

## Discussion

Research in humans has shown that individual cycle length is typically most irregular, and the variability in menstrual cycle length can range from 10 to 16 days due to varying lengths of the follicular phase (Apter et al., 1987; Bashay and Carr, 2000; Lenton et al., 1984). The estradiol peak in cynomolgus monkeys is similar to that in humans because it occurs 1 day before the peak in luteinizing hormone (LH) concentration (Shimizu, 2015). The menstrual cycle in the cynomolgus monkey consists of a 12 to 14 day follicular phase, a 3-day ovulatory phase, and a 14 to 16 day luteal phase (Weinbauer et al., 2008). Although the lengths of the ovulatory and luteal phases are consistent between cynomolgus monkeys and humans, the follicular phase is more variable in duration in cynomolgus monkeys, and as a result, affects the subsequent ovulatory phase (Esch et al., 2008; Shimizu, 2015). Thus, the length of the menstrual cycle is different for each monkey, and the timing of ovulation varies. It is unscientific to calculate an ovulation point based on the average of the menstrual cycle of all monkeys. In this study, a personalized superovulation protocol was developed based on the differences among individual menstrual cycles. It was compared with the typical superovulatory protocols based on the average of all menstrual cycles, the total numbers of oocytes retrieved and proportion of MII-stage oocytes were significantly higher than those with protocol I (26.6 ± 9.0 vs 16.4 ± 7.3 and 58.5% ± 9.9% vs 41.3% ± 25.5%, respectively; *P* < 0.05 and *P* < 0.01); and the proportion of MII-stage oocytes was higher than that with protocol II (58.5% ± 9.9% vs 34.4% ± 11.6%, *P* < 0.01). These results exceeded those observed previously for other cynomolgus monkeys (47.6% ± 32.1% MII) (Kim et al., 2017), baboons (range, 20%–43% MII) (Nyachieo et al., 2009), or rhesus monkeys (range, 44%–53% MII) (Hewitson et al., 1998; Nusser et al., 2001). The total number of retrieved oocytes and the proportion of MII-stage oocytes clearly demonstrated that the personalized superovulation protocol resulting in more mature MII-stage oocytes during the natural ovulatory cycle of mature and even older female cynomolgus monkeys (6–13 years of age). Similar to the situation in humans, each monkey’s reproductive cycle is not identical, and the ovulation timing for each individual cannot be calculated solely using the average for all monkey menstrual cycles (Ji, 2013; Yoshida et al., 1982).

Within the same monkey, some differences exist in the percentage of GV and MII-stage oocytes between the first ovulation induction using superovulation protocol I and the second ovulation induction using a personalized superovulation protocol (51.0% ± 25.9% vs 14.6% ± 11.3%, 12.8% ± 9.9% vs 51.5% ± 24.9%, respectively; *P* < 0.05 and *P* < 0.05). Similarly, the yield of oocytes in MII was greater in the second ovulation induction using the personalized superovulation protocol compared with first ovulation induction using protocol II (51.0% ± 13.2% vs 34.1% ± 2.7%, *P* < 0.05). The proportion of MII-stage oocytes showed an increasing tendency, and the proportion of GV-stage oocytes exhibited a decreasing tendency, compared with the first ovulation induction. Despite research suggesting that consecutive ovarian stimulations could reduce the quantity and quality of the oocytes obtained(Abbott et al., 2013; Lee et al., 2011; Van Blerkom and Davis, 2001), the analysis of data in the present study suggested that repeated stimulation did not impair ovarian response in terms of the quantity (17.3 ± 6.7 vs 22.5 ± 16.1, 19.3 ± 7.4 vs 17.0 ± 7.0) of oocytes obtained. Furthermore, the number of MII-stage oocytes obtained was effectively improved by adjusting the superovulation protocol to account for the menstrual cycle of each particular cynomolgus monkey. It was the fifth to sixth menses after the retrieval of oocytes from the first cycle. The cynomolgus monkeys treated with the personalized superovulation protocol had no bad influence, and the proportion of MII-stage oocytes increased, thereby allowing the effective utilization of this precious resource.

A comparison of the results of superovulation protocol II and the personalized superovulation protocol revealed that the use of GnRH-a in combination with the length of the individual menstrual cycle of cynomolgus monkey also served the role of GnRH-a in ovulation induction. GnRH-a is gonadotropin-releasing hormone agonist. The biological activity of GnRH-a is 50–200 times higher than that of endogenous GnRH(gonadotropin-releasing hormone) for the GnRH receptor. Therefore, its affinity to GnRH receptors substantially increased. It also causes LH (luteinizing hormone) and FSH (follicle-stimulating hormone) hypersecretion (flare-up) (Fouda et al., 2016; van Loenen et al., 2002). Based on this pharmacological feature, GnRH-a has been universally used for ovarian hyperstimulation cycles in human ART (Dumont et al., 2016; Zervoudis et al., 2016). After GnRH-a have induced a state of hypoestrogenism, exogenous FSH is given to stimulate ovarian follicles, in addition to synchronization with the naturally occurring FSH peak, followed by the use of hCG to trigger oocyte release (van Loenen et al., 2002).

Ovulation during the natural menstrual cycle require participation of LH/FSH peaks, in particular, the oocyte maturation process require appropriate LH peak, natural ovulation usually occur 35 to 36 h after the appearance of the endogenous LH/FSH peak(Hoff et al., 1983). The time interval of oocyte and hCG injection are varies ranges from 27-36 h for rhesus monkey, and the percentage of MII stage oocytes retrieval are 13%, 24%, 31% and 46% at intervals of 27 h, 32 h, 34 h and 36 h, respectively(P et al., 1996). Studies on cynomolgus monkeys have shown, the time interval between oocyte retrieval and hCG injection varied from 28 to 43 h, resulting in successful pregnancy (Ogonuki et al., 2003). The personalized superovulation protocol incorporated the menstrual cycle length of each cynomolgus monkey, thereby allowing to determine the timing of oocyte retrieval based on the equation: menstrual cycle length/2 ± 1, and then push this time 35-36 hours forward as the time of hCG injection, this time just close to the time of natural ovulation and consistent with the monkey’s reproductive physiological laws, and it is consistent with recent research by Liu Zhen et al (Liu et al., 2018b).

The primary focus of embryo transfer is how to ensure the synchronization of the recipients, currently, there are two methods most commonly used. The first method is to detect the concentration of estradiol (E2), the rapid decline in the plasma steroid hormone E2 peak (usually below 100 pg/mL) that can predict the time of ovulation, usually 2-3 days after the E2 peak occurs, the corresponding embryo can be transferred into the oviduct or uterus. Some studies have used this method to determine the synchrony of recipients and obtain offspring(Niu et al., 2010; Sun et al., 2008), however, there are three problems with this method: 1. All selected monkeys begin venous blood collection on the fourth day of menstruation and test E2 daily for about 8 to 19 days, we have also used this method to determine the synchrony of surrogate rhesus monkey before, but this process is cumbersome, and daily blood collection will cause stress injury and blood loss to the monkeys, which will have an adverse effect on implantation of embryos. 2. Stigma of ovulation on the ovary may not be visible on the 2-3^th^ day after the E2 peak appears, however, an important feature of recipients synchronization is the formation of stigma of ovulation or corpus luteum. 3. According to previous test data, this peak occurs at a wide range from 7 to 20 days after menstruation (Fig. 4A), and there are obvious individual differences, this means that the same batch of embryos can be transferred on the 8^th^ day of menstruation, but some need to be cultured in vitro for up to 13 days before they can be transferred. Cynomolgus monkey embryos can develop to blastocysts in vitro culture for 7 days (about 168 hours), in vitro culture for up to 13 days will inevitably greatly reduce the quality of the embryo, and at this time, the developmental state of the embryo is suitable for transferring into uterus of the recipient. However, due to the hard and narrow curvature of the cervix monkey, the oscilloscope has a dome structure, uterine transplantation is not conducive to the passage of the embryo transfer tube, and the actual operation is very difficult, embryo transfer lower implantation rate(Wolf, 2004; Wolf et al., 2004), so the uterus transplantation has basically been abandoned in non-human primates. The second method of ensure the synchronization of the recipients is that the female recipients ovaries had a stigma, and the surrogate mother of the world’s first somatic cell clone monkey is also obtained by this method (Liu et al., 2018b). This method is highly reliable for ensuring synchronization. However, in previous studies, we found that monkeys have normal menstrual cycles, but no stigma. It is speculated that this phenomenon may be similar to humans, in humans called anovulatory menstruation. When the follicle develops only to a certain extent, but is immature, it can secrete a certain amount of estrogen, however, the peak of estrogen secretion is not reached, and the ovary can not ovulate. No stigma formation, there is no secretion of progesterone, so that the endometrium has been in the proliferative phase and can not be converted into a secretory phase, although the uterus bleeding on time, but not true menstruation, but anovulatory menstruation(Hambridge et al., 2013; Mumford et al., 2011). Therefore, relying on the monkey’s menstrual cycle to go directly to the stigma, there is uncertainty, may reduce the efficiency of obtaining a surrogate mother.

In summary, in order to avoid the above shortcomings, we fully considered the possible factors affecting ovulation according to the reproductive physiology characteristics of non-human primates, based on 3 points to ensure the recipients synchronization efficiently and accurately, that is, “the normal menstrual cycle, sexual swelling and stigma of ovary”. Sexual skin characteristics of young female monkey of the early stage of sexual maturity and adult female are different, see method and Fig. 4C and 4E for specific characteristics. Overall, during the high estrus period, due to the influence of estrogen, abnormal engorgement occurs in the external genital area of female cynomolgus monkeys. The color of this sexual skin becomes darker (mostly red), and swelling was a phenomenon that we call "sexual swelling" (Gillin and Birrell, 1996). Normal "sexual swelling" occurs during ovulation, which is on day 13-19 after menstruation onset. After observing the change in the color of sexual skin of surrogate candidate, more than 98% of monkeys with normal menstrual cycles can clearly see the stigma of ovulation on the ovary by laparoscopy. From the observation of “sexual swelling”, the window period for embryo transfer can last 6-7 days. Christian R. et al. also clearly pointed out that the sex skin was very reliable way of ovulation in monkeys (Gillin and Birrell, 1996).

Based on the previous personalized ovulation induction and comprehensive method of selecting recipients, we obtained 2 cases of successful pregnancy from a high-quality 2-4-cell embryos that were obtained from a donor for transplantation into the oviduct of 4 surrogate mothers, as a result, 3 fetuses were aborted on day 103. This does not necessarily mean that offspring can be obtained easily by ICSI, because ours was the first case in which a monkey had 3 fetuses based on the personalized superovulation in nonhuman primate. This study concluded that the personalized superovulation protocol increased the retrieval rate of MII-stage oocytes, and through the ICSI, the embryos were successfully obtained and transplanted into the oviducts of the surrogate mothers selected according to the “menstrual cycles”, “sexual swelling”, “stigma on the ovary”, and the pregnancy was successful. Which proved that our method of ensuring surrogate mother synchronization was not only simple and reliable but scientific and efficient. Thus enhanced the efficiency of ART in cynomolgus monkeys and improved the success of generating genetically modified monkeys under the premise of minimizing damage to animals.

## MATERIALS AND METHODS

### Animals

Twenty-four adult female cynomolgus monkeys (*Macaca fascicularis*; 6–13 years of age; mean, 9.04 years) were selected for hormonal stimulation and were used for oocyte collection, additional four adult females (4.5–9 years of age; mean, 7.5 years) were selected for embryo transfer, with menstrual cycles between 28 and 33 days in length, and body weights between 3 and 6 kg, and there was another sexually mature male monkey selected to collect sperm, these monkeys were provided by the Institute of Medical Biology, Chinese Academy of Medical Science, and Peking Union Medical College. The design and procedures of all animal studies were in accordance with the ethical standards detailed in the 1964 Declaration of Helsinki and its later amendments. All animal works were approved by the Yunnan Province Experimental Animal Management Association and the Experimental Animal Ethics Committee of the Institute of Medical Biology Chinese Academy of Medical Sciences which based on the 3R principle (reduction, replacement, and refinement).

### Superovulation protocols

We investigated 3 protocols for superovulation of female cynomolgus monkeys. Superovulation protocol I: Female cynomolgus monkeys that had not previously received ovarian stimulation were administered single intramuscular injection of 25 IU follicle-stimulating hormone (FSH, Livzon Pharmaceuticals Co., Ltd, China) on day 3 of the menstrual cycle. On the fourth day of the menstrual cycle, they were injected twice daily with FSH for 8 days, followed by an intramuscular injection of chorionic gonadotropin (hCG, 1500 IU, Livzon Pharmaceuticals Co., Ltd) on the 12^th^ day of the menstrual cycle, 35–36 h after collecting oocytes [i.e., oocyte harvest was on the 13^th^ day of the menstrual cycle (Fig. 2A)]. Superovulation protocol II: female cynomolgus monkeys that had not previously received ovarian stimulation, the difference from superovulation protocol I being that a subcutaneous injection of 0.1 mg of gonadotropin-releasing hormone (GnRH-a, Ferring Pharmaceuticals Ltd, Germany) on the first day of the menstrual cycle (Fig. 2B). Personalized superovulation protocol : regardless of whether female cynomolgus monkeys had previous ovulation induction experience, they were given a subcutaneous injection of 0.1 mg GnRH-a on the first day of the menstrual cycle. After a day’s interval, on the third day of the menstrual cycle, an intramuscular injection of FSH (25 IU) once a day was administered starting on day 4 of the menstrual cycle and then twice daily for 8–12 days. The final FSH dose (25 IU) was administered once daily on the 12–15^th^ days based upon the length of each monkey’s menstrual cycle. If the menstrual cycle was 28 days, hCG (1500 IU) was injected on the 12^th^ or 13^th^ day, with oocyte harvest 35–36 h later (i.e., on the 14^th^ or 15^th^ day of the menstrual cycle). If the menstrual cycle was 30 days, hCG (1500 IU) was injected on day 13 or 14, followed by oocyte collection 35–36 h later (on the 15^th^ or 16^th^ day of menstrual cycle). The time of oocyte retrieval was based on the principle of menstrual cycle length/2 ± 1, and the latest date for harvesting oocytes was on day 17 of the menstrual cycle (Fig. 2C).

### Oocyte retrieval and in vitro culture

For oocyte retrieval (usually 35–36 h after hCG injection), all female monkeys were anesthetized with 3% pentobarbital sodium at a dose of 25 mg per kg of body weight. Ovaries were exposed through a small incision in the middle of the lower abdomen, and the contents of large follicles (1 cm ± 5 mm in diameter) were aspirated through a 25-gauge needle connected to a 5.0-mL syringe. Cumulus oocyte complexes (COCs) were harvested by aspiration with the medium [TALP-HEPES (Tyrode’s Albumin Lactate Pyruvate, Caisson Labs, USA), supplemented with 5 mg/mL bovine serum albumin (Sigma-Aldrich, USA) and 5 IU/mL of heparin (Sigma-Aldrich, USA)] under a stereomicroscope (Nikon SMZ745, JAPAN). COCs were rinsed with TALP-HEPES supplemented with 0.1% hyaluronidase (Sigma-Aldrich, USA) for removing cumulus cells. The maturation status of the oocytes was evaluated under an inverted microscope (Leica DMI6000B, Germany) at ×100 or × 200 magnification, and GV, MI, and MII oocytes were counted for statistical analysis after washing. Oocytes (both immature and mature) were cultured in 50-uL drops of defined medium [HECM9 medium (Gibco, Invitrogen, Gand Island, USA), containing 10% FBS ((Sigma-Aldrich, USA) and 10% GLU ((Sigma-Aldrich, USA)] under a layer of embryo-tested mineral oil (Sigma-Aldrich, USA)) at 37°C in 5% CO_2_ and 95% compressed air at high humidity (Thermo 3131, Waltham MA, USA).

### Sperm collection

Sperm was obtained by electric stimulation of adult male cynomolgus monkey penis. Washed it twice times through TALP-HEPES supplemented with 5 mg/mL bovine serum albumin, centrifuged at 2000 rpm for 5min and discard the supernatant. The last time added 1.0 mL TALP-HEPES supplemented with 5 mg/mL bovine serum albumin, waiting for 5 minutes for ICSI, that is, used the upstream method to take the sperm with good viability in the supernate.

### Intracytoplasmic sperm injection

The MII-stage oocytes derived from a monkey that was induced by personalized superovulation protocol were cultured at least 2 h before ICSI. ICSI was conducted using a inverted microscope (Leica Microsystems, LEICA-DMI6000B) with attached micromanipulators, briefly, separately placed mature oocytes and sperms into two 10 μL droplets of TALP-HEPES containing 5 mg/mL bovine serum albumin, and the motile spermatozoa were immobilized in 7% polyvinylpyrrolidone (PVP). The oocyte was fixed in holding pipette, and the first polar body was placed in 12 point position, the injection pipette penetrated oocytes at 3 o’clock, passed through the zona pellucida and oocyte membrane respectively, negative pressure to attract lightly broken membrane after the sperm slowly injected into the oocyte, and then gently backed off injection pipette. Finished the sperm injection of all MII-stage oocytes and transferred the oocytes to HECM9 medium and incubated at 37°C in 5% CO2 and 95% compressed air.

### Recipients selection and embryo transfer

Healthy, sexually-mature and normal menstrual cycle (mean, 28–33 days) female cynomolgus monkeys (age 4.5–9 years, body weight 3.5–5 kg) were selected as embryo recipients. And it is normal for these monkeys to be observed at least 3 times or more of their menstrual cycle before they can be used as a surrogate mother. At the same time, accurate observation and judgment of “sexual swelling”. Observation of sexual skin of young female monkeys: young female in the early stage of sexual maturity, the vaginal area is swollen, reddened and subcutaneous groin subcutaneously forms two saclike protrusions of testicular size as the main features, and it has a smooth, shiny appearance and is deep, intense red color, periodic changes, and then did not appear (Fig. 4C); Observation of sexual skin of adult female monkeys: turgescent area extended to anus and thighs, the perineal skin is fully distended with no wrinkles and smooth and most intense bright red color (Fig. 4E). Normal "sexual swelling" occurs during ovulation, which was on day 13-19 after menstruation onset. After observing the change in the color of sexual skin of surrogate candidates, checked their ovaries where have a stigma of ovulation via laparoscopy. From the observation of “sexual swelling”, the window period for embryo transfer could last 6-7 days.

### Parentage verification of aborted fetus

Parentage verification of the aborted fetus were done by DNA typing of 20 microsatellite markers: 3 fetuses took the hind limbs to extract DNA, oocytes donor, sperm donor and surrogate mothers took blood sample to extract DNA. 20 locus-specific primers were designed according to references (Kanthaswamy et al., 2010; Siddiqui et al., 2009) and each primers contained a fluorescent dye (FAM/HEX) (Table 1), PCR amplification of STRs was performed as follows: firstly, pre-denaturation at 95°C for 1 minute, then was amplified with 35 cycles of 94°C for 30 s, 56°C for 30 s, and 72°C for 30 s, followed by a 30-min extension at 60°C. The multiplexed reaction containing 10×Buffer I (Beijing Microread), 2.5mMdNTP, –F+R (5Ìm FAM or HEX –labeled)(Takara, JAPAN), HSTaq (Takara, JAPAN) and DNA template. PCR were carried out in ABI GeneAmp9600. Then, add a mixture of internal size standard ORG 500(Beijing Microread), deionized formamide (0.5:8.5) and PCR product to a 96-well plate. Followed by capillary electrophoresis on ABI 3730XL DNA analyzer to obtain the raw data. The raw data file detected by the 3730XL was imported into the analysis software genemapper ID3.2 for analysis. According to genetic principles and genetic laws, half of the alleles in the progeny genotypes are from the male parent and the other half are from the female parent.

**Table 1.** The 20 STRs and Primers Used in This Study.

### Statistical analysis

Oocytes data were analyzed using GraphPad Prism 5.0 (GraphPad Software, free trial version, California San Diego, USA). All data were expressed as the means ± standard deviation. Comparisons of the number and percentage of oocytes obtained using the three protocols were performed using the nonparametric Kruskal–Wallis test. The results were considered highly significantly different if *P* ≤0.01, and significantly different if *P* ≤0.05.

## Acknowledgements

We thank Jiahong Gao and Cong Li (Institute of Medical Biology, Chinese Academy of Medical Sciences and Peking Union Medical College) for their technical assistance.

## Competing interests

The authors declare no competing or financial interests.

## Author contributions

Kaili Ma and Qihan Li designed the experiments; Zhangqiong Huang performed embryo culture, ICSI and embryo transfer. Yun Li performed ovulation induction, recipients selection and sperm collection. Qinfang Jiang, Yixuan Wang conducted oocytes retrieval and garentage verification of aborted fetus, Kaili Ma, Qihan Li and Zhangqiong Huang interpreted the data and wrote the manuscript. All authors prepared the figures and approved the final version of the manuscript.

## Funding

This work was supported by the National Natural Science Foundation of China (81571254,81301073); the Joint Funds of the National Natural Science Foundation of China (U1402221); National Science and Technology Major Project of the Ministry of Science and Technology of China (2016ZX08011007-003); Yunnan Natural Science Foundation (2016FA032); the Fundamental Research Funds for the Central Universities (3332016116,2016ZX310179-1); CAMS Innovation Fund for Medical Sciences (2016-I2M-2-001,2016-I2M-1-0040); the Foundation for the Institute of Pathogen Biology (2015IPB201) and the Foundation for the Institute of Medical Biology (2014IMB01ZD)

## Data availability

No data availability

---

**Data 1** Extensive Lists of STRs Examined for Aborted Fetus a, b and c.

